# Stratigraphic distribution of ammonoids and other macrofossils from the upper Albian–lower Cenomanian interval in Putumayo, Colombia, northern South America: new data for mid-Cretaceous basin correlation

**DOI:** 10.1101/2023.04.24.536408

**Authors:** Javier Luque, Jonatan Bustos, Alejandro Beltrán-Triviño, Angie Rodriguez, Alexandra Delgado, Johan M. Sanchez, Julián Caraballo, Manuel Paez-Reyes, Mikel A. López-Horgue

## Abstract

Despite the rich paleontological heritage of Colombia, in the equatorial Neotropics, one of the least explored regions in terms of its fossil record is the Putumayo region near Ecuador, largely due to its considerable ground cover, thick vegetation, rock weathering, geographic remoteness, and overall inaccessibility to well-exposed outcrops. This precludes detailed comparisons with neighboring basins, and thus the generation of more comprehensive biostratigraphic correlations for western northern South America and other paleobiogeographic regions, e.g., Mediterranean Tethys, northern Africa, Western Interior Basin. Here, we report 67 occurrences of mid-Cretaceous ammonoids and other macrofossils (e.g., bivalves, decapod crustaceans, fish remains, plant remains), from the middle Albian of the uppermost Caballos Formation and the upper Albian–lower Cenomanian of the lower Villeta Formation, collected in-situ from a stratigraphic section cropping out on the Mocoa–San Francisco road in the Department of Putumayo, Colombia. Among the ammonoid taxa recovered are several morphotypes assignable to ?*Schloenbachia* sp., cf. *Engonoceras* sp., *Oxytropidoceras* (*Venezoliceras*) sp., *Oxytropidoceras* (*Laraiceras*) sp., cf. *Oxytropidoceras* sp., *Mortoniceras* (*Mortoniceras*) cf. *vespertinum*, *Mortoniceras* (*Mortoniceras*) sp., *Algericeras* sp., *Hysteroceras* sp. 1, *Hysteroceras* sp. 2, cf, *Hysteroceras* sp., cf. *Forbesiceras* sp., *Graysonites* sp., *Hamites* sp., and two isolated aptychi. The occurrence of the ammonoid genera *Oxytropidoceras*, *Mortoniceras*, and *Schloenbachia*, suggest interconnection of the Putumayo Basin during the mid-Cretaceous with the Upper Magdalena Valley in Colombia and the Oriente Basin in Ecuador, which, together with the rest of the ammonoid assemblage, provide biostratigraphic data to define the upper Albian–lower Cenomanian in the basin and thus in northwestern South America.

## 1. Introduction

The Putumayo region, located in southwestern Colombia near the border with Ecuador and Perú, is key to understanding the regional geological evolution of northern South America. Unfortunately, despite its rich geological and paleontological record and oil and gas production, due to its geographic, topographic, and biological characteristics, it is one of the least studied areas compared to neighboring basins, e.g., the Middle Magdalena Valley Basin, Upper Magdalena Valley Basin, and the Llanos Basin (Colombia), the Oriente Basin (Ecuador), and the Marañón Basin (Perú) (<u>Saeid et al., 2017</u>), and thus with meager knowledge of its paleontology and biostratigraphy.

Among the few paleontological studies in the area are the reports of <u>Royo y Gómez (1942)</u> and <u>Cucalón and Camacho (1966)</u>, who mentioned the occurrences of “Paleohoplitidae”, *Oxytropidoceras multifidum*, *Neoplycticeras* (?) *subtuberculatum*, *Neophlycticeras rhombifera*, *Mortoniceras*, *Neophyeticeras*, *Corbula*, *Astarte*, *Pecten compressus*, *Inoceramus plicatus*, *Inoceramus labiatus* and *Exogyra squamata*, but unfortunately, illustrations of these specimens and their stratigraphic context are lacking. Moreover, the transition between the upper Albian and the lower Cenomanian in Colombia and in northern South America is poorly understood, and the contact between both ages has proven to be elusive (<u>Renz, 1982</u>).

Here, we report new occurrences of uppermost Lower Cretaceous and lowermost Upper Cretaceous ammonoids, bivalves, decapod crustaceans, and other macrofossils from Putumayo, Colombia, collected *in-situ* from an Albian–lower Cenomanian stratigraphic section cropping out along the Mocoa–San Francisco road (hereafter referred to as San Francisco) (Figs 1 and 2). Ammonoid faunas from San Francisco provides further evidence to support the presence of the Albian–Cenomanian transition in the northern Andes, as indicated by the rapid faunal turnover from several genera circumscribed to the Albian (e.g., *Oxytropidoceras*) to genera that have their first occurrence in the Cenomanian (e.g., *Schloenbachia*, *Forbesiceras*, *Graysonites*, *Algericeras*) (<u>Young, 1958</u>; <u>Kennedy and Wright, 1981</u>; <u>Kennedy and Klinger, 2008</u>; <u>Kennedy and Klinger, 2011</u>; <u>Kennedy, 2013</u>) (Figs 2–7). In this work, we contribute to the biostratigraphy of ammonoids across the upper Albian–lower Cenomanian in the equatorial Neotropics, filling the gap in our understanding of the geological and paleontological history of the Putumayo Basin and therefore on the oil source rocks in the basin, and discuss their implications to recognize the Lower Cretaceous–Upper Cretaceous boundary in northwestern South America.

**Figure 1.**
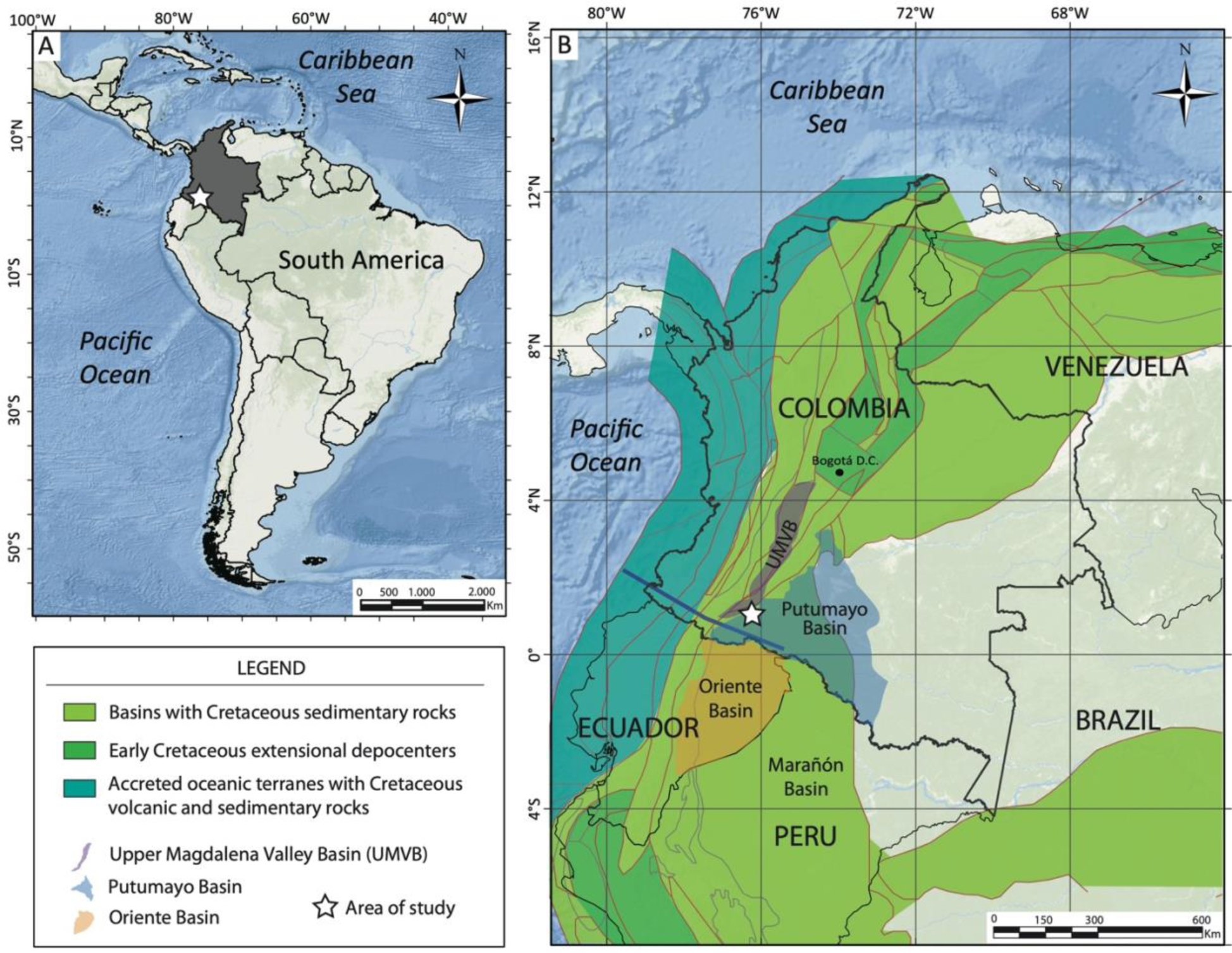
Location map of the study area. **A**, General map of South America. **B**, General map of Colombia, showing the overall distribution of Cretaceous-only rocks and depocenters. White star indicates the studied Albian–Cenomanian section in the Mocoa–San Francisco section, Putumayo, Colombia, bearing fossil macroinvertebrates. Map in B modified after <u>Sarmiento-Rojas (2019)</u>.

**Figure 2.**
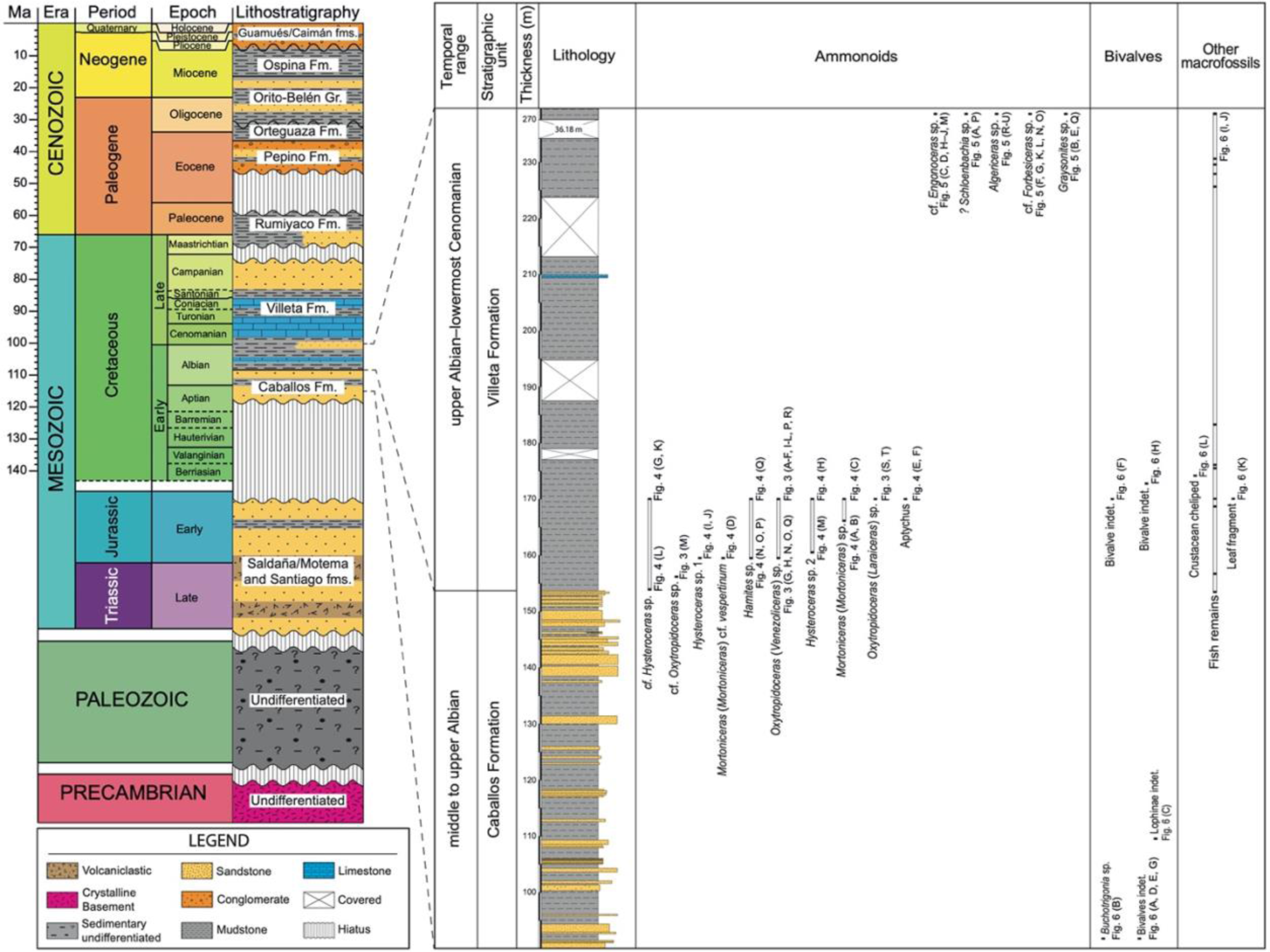
Litho-and biostratigraphy of the mid-Cretaceous in the Putumayo region, Colombia, northern South America. Left: Simplified, generalized lithostratigraphic column of the units in the area of study. Center: Lithostratigraphic column of the studied upper Albian–lower Cenomanian interval of the Villeta Formation in the Mocoa–San Francisco section (this study). Right: Stratigraphic ranges of ammonoids and other macrofossil occurrences (e.g., bivalves, crustacean remains, fish remains, etc.). A full, expanded, detailed stratigraphic column is included as Supplementary Appendix 1.

**Figure 3.**
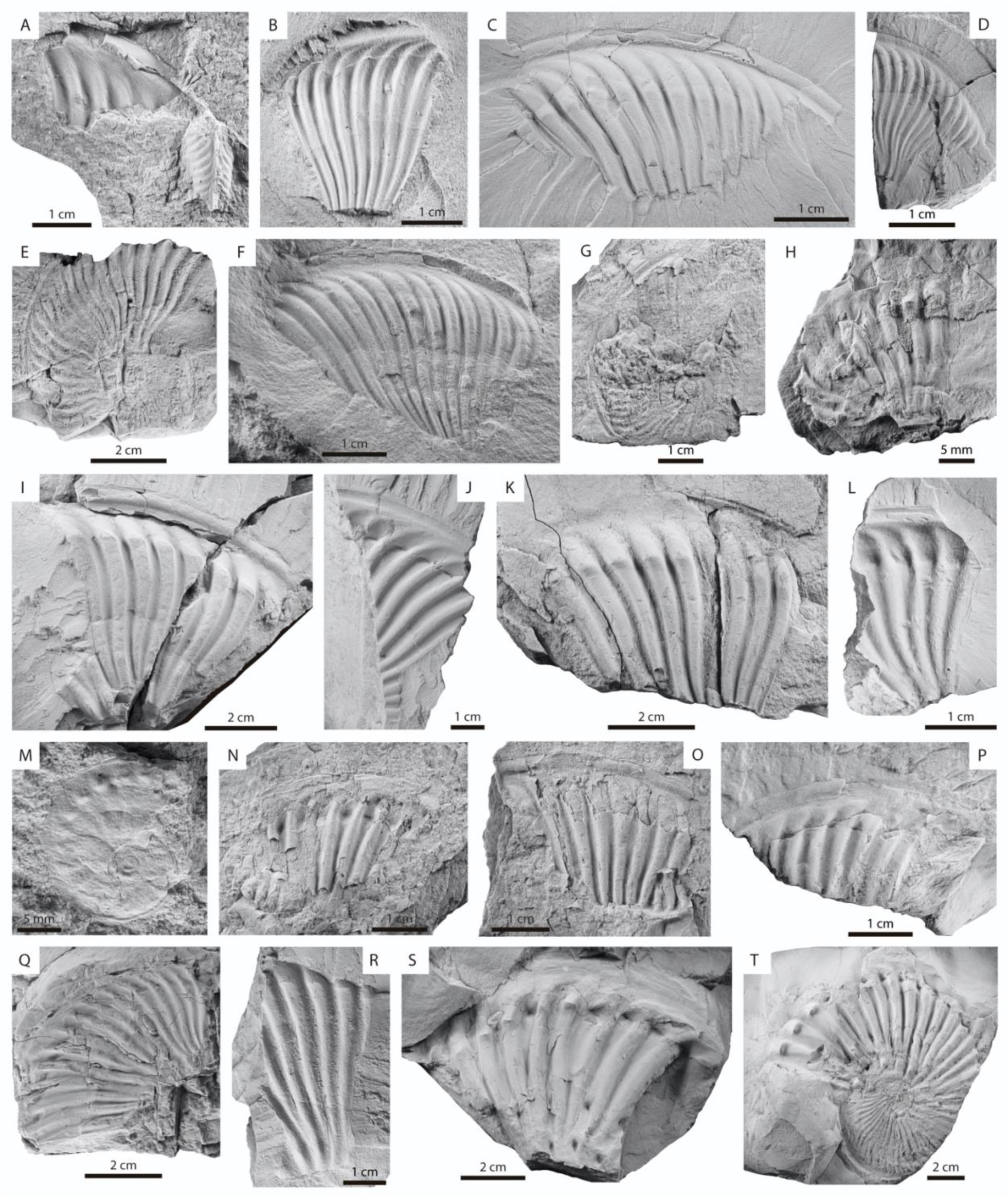
Upper Albian ammonoids of the Villeta Formation, Putumayo, Colombia. **A–L**, **N–R**, *Oxytropidoceras* (*Venezoliceras*) sp. **A**, **B**, **D**, **F**, specimens showing a keel. **A**, AGESS-P-0169-03, two individuals, the smallest one showing the ventral view (bottom right), the largest one is an external mold in lateral view (upper left). **B**, AGESS-P-0156-12, lateral view of the left side. **D**, AGESS-P-0156-10, lateral view of the right side. **F**, AGESS-P-0169-04, lateral view of the right side. **E**, **G**, most complete specimens, showing the involute coiling of the last preserved whorl. **E**, AGESS-P-0156-13, lateral view of the left side. **G**, AGESS- P-0166-05, external mold in lateral view of the right side. **C**, **H–L**, **N–R**, specimens showing a keel and ribs bearing ventrolateral clavi. **C**, AGESS-P-0156-17, lateral view of the right side. **H**, AGESS-P-0166-10, lateral view of the left side. **I**, AGESS-P-0156-06, lateral view of the right side. **J**, counterpart of **I**. **K**, AGESS-P- 0156-07, lateral view of the right side. **L**, AGESS-P-0156-18, lateral view of the right side. **O**, AGESS-P-0166- 16, lateral view of the right side. **N**, counterpart of **O**. **P**, AGESS-P-0169-06, lateral view of the right side. **Q**, AGESS-P-0166-09, external mold in lateral view of the right side. **M**, cf. *Oxytropidoceras* sp., AGESS-P-0159-08, small specimen showing the involute coiling of the last preserved whorl and ventrolateral tubercles. **S**, **T**, *Oxytropidoceras* (*Laraiceras*) sp., AGESS-P-0156-01, showing a keel and three rows of tubercles (umbilical, low lateral, and ventrolateral) bearing on the long ribs, the ventrolateral tubercle has a clavus-like shape in its inner and outer terminations; **T**, lateral view of the right side; **S**, counterpart of T. All specimens imaged dry and coated with ammonium chloride.

**Figure 4.**
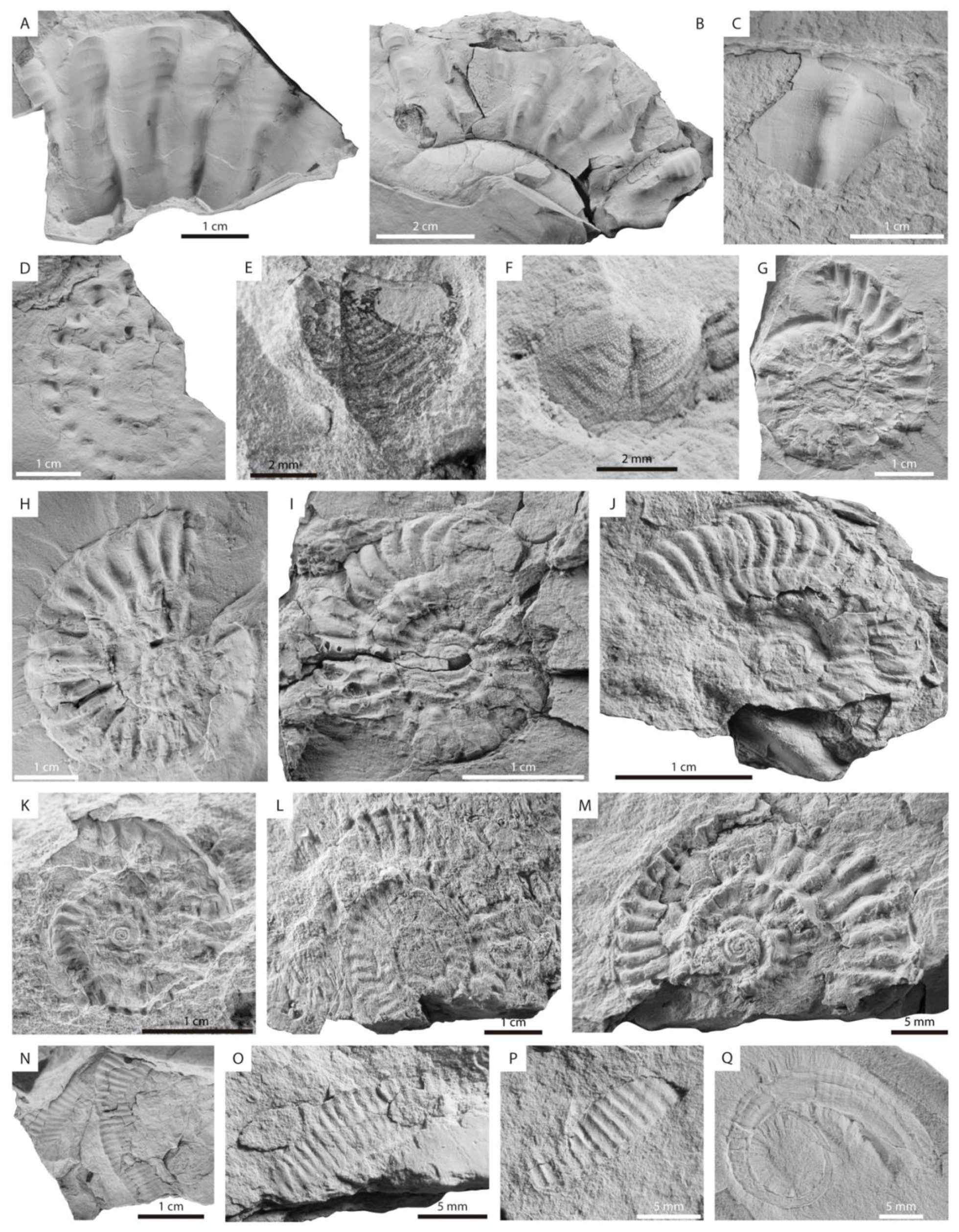
Upper Albian ammonoids of the Villeta Formation, Putumayo, Colombia (cont.). **A–C**, *Mortoniceras* (*Mortoniceras*) sp. **A**, **B**, AGESS-P-0168, specimen showing trituberculate ribs; **A**, counterpart of B; **B**, lateral view of the right side. **C**, AGESS-P-0169-02, external mold in lateral view of the right side. D, AGESS-P- 0166-04, *Mortoniceras* cf. *vespertinum,* external mold in lateral view of the right side showing two rows of tubercle. **E**, **F**, aptichi showing articulate preservation of the two halves; **E**, AGESS-P-0156-22; **F**, AGESS-P- 0156-23. **I**, **J**, *Hysteroceras* sp. 1, small specimens showing evolute coiling of the last preserved whorl. **I**, AGESS-P-0166-17, lateral view of the left side. **J**, AGESS-P-0166-12, lateral view of the right side. **H**, **M**, *Hysteroceras* sp. 2, small specimens showing evolute coiling of the last preserved whorl. **H**, AGESS-P-0156- 04, external mold in lateral view of the right side. **M**, AGESS-P-0167-01, external mold in lateral view of the right side. **G**, **K**, L, cf. *Hysteroceras* sp. small specimens showing evolute coiling of the last preserved whorl; **G**, AGESS-P-0156-19, external mold in lateral view of the left side; **K**, AGESS-P-0156-21, external mold in lateral view of the left side; L, AGESS-P-0158-06, external mold in lateral view of the left side. **N–Q**, *Hamites* sp. **N–P**, fragmentary specimens showing single radial ribs. **N**, AGESS-P-0166-01, external mold; **O**, AGESS- P-0166-02, external mold; **P**, AGESS-P-0166-03, external mold; **Q**, AGESS-P-0156-02, early whorls showing a crioconic coiling and fine radial ribs. All specimens imaged dry and coated with ammonium chloride.

**Figure 5.**
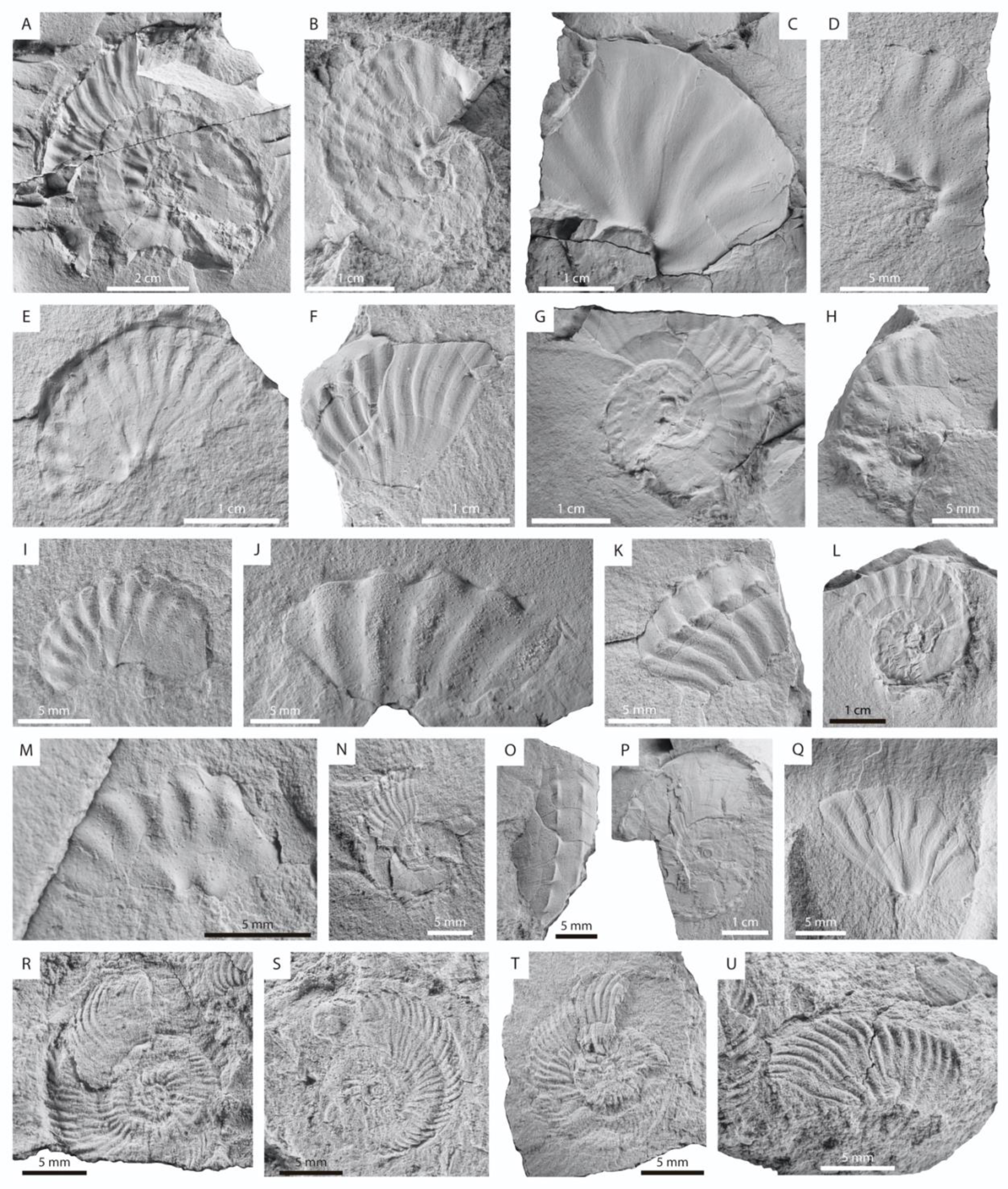
Lower Cenomanian ammonoids of the Villeta Formation, Putumayo, Colombia. **A**, **P**,?*Schloenbachia* sp. **A**, AGESS-P-0155-01, external mold in lateral view of the right side, showing coiling of the last preserved whorl and umbilical tubercles. **P**, AGESS-P-0155-12, lateral view of the left side, showing fine ribbing and ventrolateral clavi. **C**, **D**, **H**–**J**, **M**, cf. *Engonoceras* sp. showing slightly conical umbilical tubercles and wide flexuous ribs. **C**, AGESS-P-0155-03, lateral view of the right side. **D**, AGESS-P-0155-08, lateral view of the right side. **H**, AGESS-P-0155-14, lateral view of the left side. **I**, AGESS-P-0155-15, lateral view of the left side. **J**, AGESS-P-0155-21, lateral view of the right side. **M**, AGESS-P-0155-11, lateral view of the right side. **F**, **G**, **K**, **L**, **N**, **O**, cf. *Forbesiceras* sp., with F, K, O, showing ribs bearing ventrolateral clavi. **F**, AGESS-P-0155-09, lateral view of the left side. **K**, AGESS-P-0155-04, oblique view showing the upper part of the flank and the ventral area. **O**, AGESS-P-0155-22, ventral view. **G**, **L**, **N**, showing coiling of the last preserved whorl. **G**, AGESS-P0155-10, lateral view of the right side. **L**, AGESS-P-0155-05, lateral view of the right side. **N**, AGESS-P0155-07, lateral view of the left side. **B**, **E**, **Q**, *Graysonites* sp. showing three rows of tubercles (umbilical, inner- and outer- ventrolateral). **B**, AGESS-P-0155-02, external mold in lateral view of the right side. **E**, AGESS-P-0155-06, external mold in lateral view of the right side. Q, AGESS-P-0155-24, external mold in lateral view of the left side. **R–U**, *Algericeras* sp.; small specimens showing numerous fine ribs strongly project in the ventrolateral region. **R**, AGESS-P-0155-17, external mold in lateral view of the right side. **S**, AGESS-P-0155-18, external mold in lateral view of the left side. **T**, AGESS-P-0155-19, lateral view of the left side. **U**, AGESS-P-0155-20, external mold in lateral view of the left side. All specimens imaged dry and coated with ammonium chloride.

**Figure 6.**
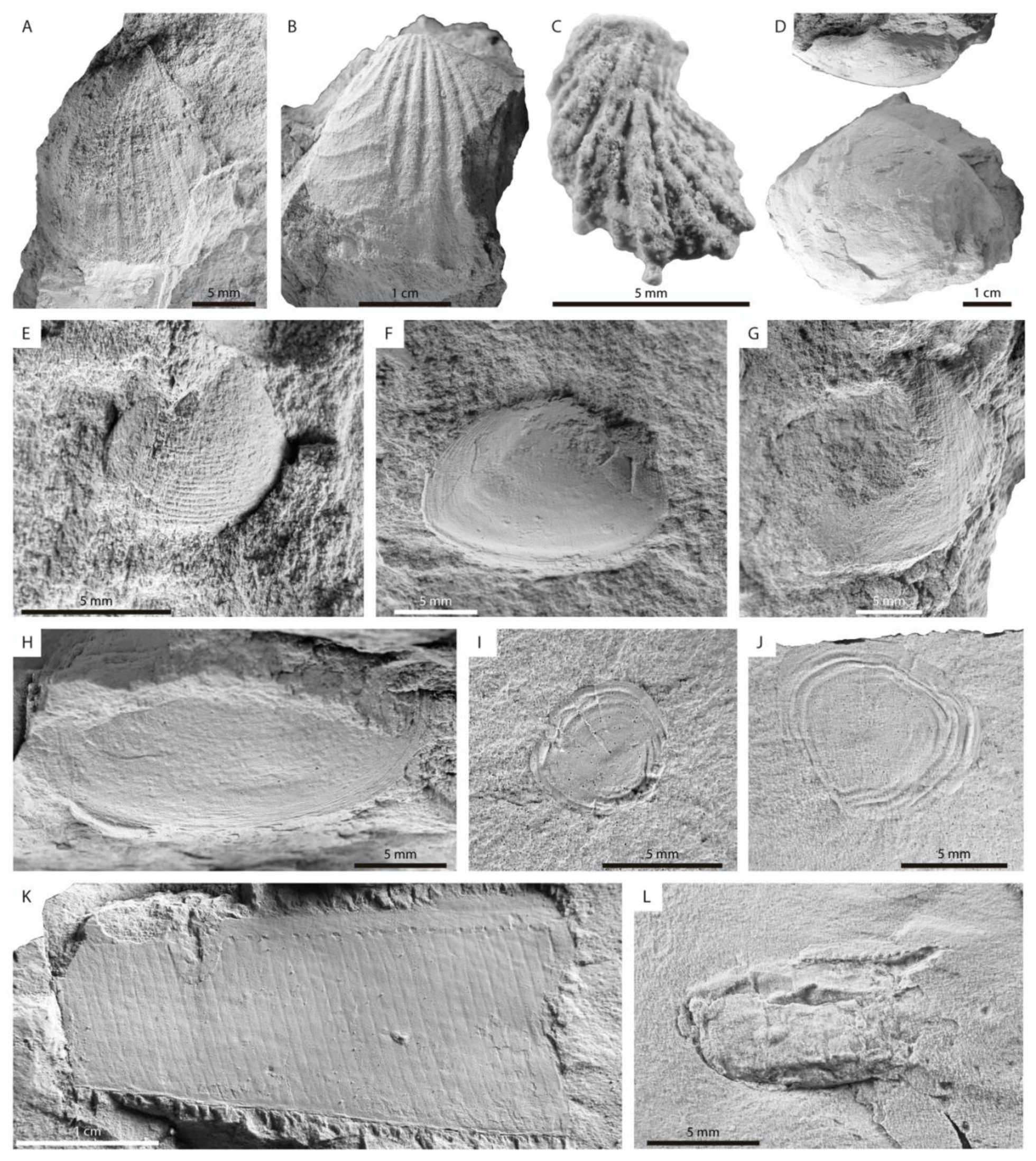
Middle Albian, upper Albian, and lower Cenomanian macrofossils of the Caballo and Villeta formations in the San Francisco section, Putumayo, Colombia. **A–E**, **G**, ?middle Albian bivalves of the Caballos Formation. **B**, *Buchotrigonia* sp. AGESS-P-0175-04, left valve. **A**, AGESS-P-0175-05, bivalve indet. **C**, AGESS-P-0161, Lophinae indet. **D**, AGESS-P-0175-06, bivalve indet. **E**, AGESS-P-0175-02, bivalve indet. **G**, AGESS-P-0175-03, bivalve indet. **F**, **H**, **K**, **L,** upper Albian macrofossils of the Villeta Formation. **F**, AGESS-P-0169-05, bivalve indet. **H**, AGESS-P-0170-03, bivalve indet. **K**, AGESS-P-0169-01, leaf fragment. **L**, AGESS-P-0170-02, decapod crustacean propodus of cheliped, possibly of a callianassid shrimp. **I**, **J**, lower Cenomanian fish scales of the Villeta Formation; **I**, AGESS-P-0155-26; **J**, AGESS-P-0155-25. All specimens imaged dry and coated with ammonium chloride.

**Figure 7.**
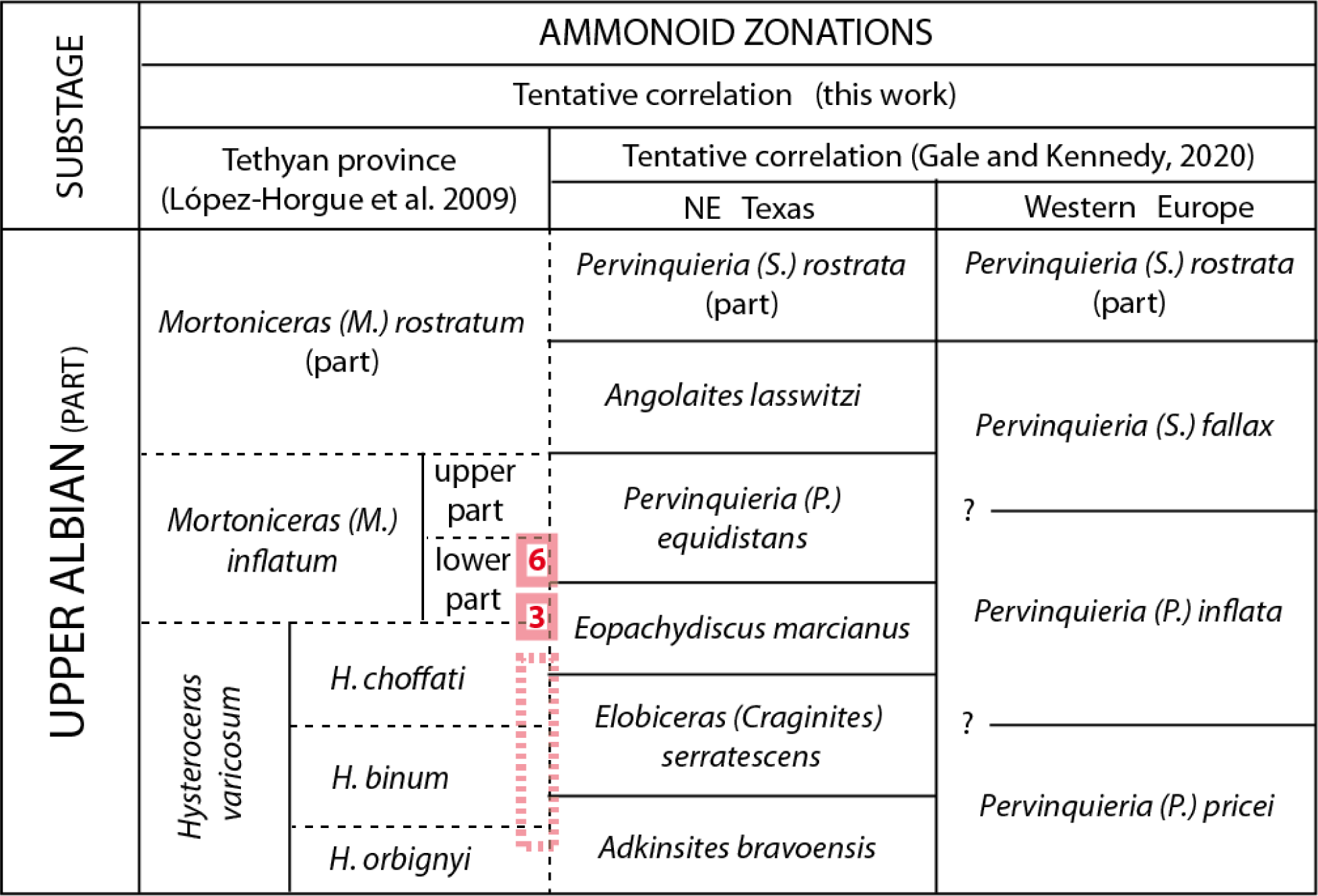
Biostratigraphic correlation of the upper Albian ammonites of the Putumayo area with the zonation of the Tethyan province, NE Texas, and Western Europe (<u>López-Horgue et al., 2009</u>; <u>Gale and Kennedy, 2020</u>). The association *Mortoniceras* (*Mortoniceras*) cf. *vespertinum*, *Oxytropidoceras* (*Venezoliceras*) sp. and *Hysteroceras* sp. 1 of level 3 (in red) is located in the transition of the *Hysteroceras choffati* to *Mortoniceras* (*Mortoniceras*) *inflatum* Zones. The association of the *Mortoniceras* (*M*.) sp. and *Hysteroceras* spp. in level 6 (in red) is probably in the lower *Mortoniceras* (*M*.) *inflatum* Zone. Dotted pink rectangle indicates the likely age for levels 1 and 2.

## 2. Geological Setting

The Putumayo Basin (Fig. 1) is a foreland basin in southern Colombia, which is considered as the northern extension of the Marañon (Perú) and Oriente (Ecuador) basins, located at the eastern flank of the Andean Cordillera (e.g., <u>Sarmiento-Rojas, 2019</u>). The stratigraphy of the Putumayo Basin can be first described by three major tectono-stratigraphic cycles separated by regional unconformities: i.e., the Pre-Cretaceous, Cretaceous, and Cenozoic cycles (e.g., <u>Gonçalves et al.,</u> <u>2002</u>). Pre-Cretaceous rocks consist of an Upper Triassic–Lower Jurassic volcano sedimentary sequence (Motema Formation, Fig. 2), which is overlaid by the uppermost Lower Cretaceous (Aptian–Albian) clastic deposits of the Caballos Formation (Fig. 2). The Cretaceous sedimentary cycle concluded with a maximum transgression, represented by the Late Albian to Santonian clastic/calcareous succession informally referred in the literature as the Villeta Formation (Fig. 2). The transgressive sequence ended during the Campanian, and then took place the subsequent progradation of continental and transitional deposits during the Late Maastrichtian and Early Paleocene (Arena N and Rumiyaco formations) (Fig. 2). In the Eocene, the deposition in the basin was dominated by fluvial/alluvial sandstones and conglomerates (Pepino Formation, Fig. 2), followed by the deposition during the Oligocene of lacustrine/fluvial shales, siltstones, and sandstones of the Orteguaza Formation (Fig. 2). Later, the continental to transitional shales, sandstones and conglomerates of the Orito-Belén and the Ospina formations were deposited since late Oligocene to early Pliocene. Ultimately, the area was covered by the coarse siliciclastic deposits of the Guamués and Caimán formations during the Pliocene and Quaternary (Fig. 2).

The sedimentary infill of the Putumayo Basin was controlled by the tectonic evolution since the Late Triassic–Early Jurassic, when the Motema Formation filled grabens formed in a back-arc setting (<u>Casero et al., 1997</u>). Later on, in the Aptian–Maastrichtian interval, the Caballos and Villeta formations accumulated in a broad and shallow epicontinental sea that extended throughout most of northwestern South America (e.g., <u>Gonçalves et al., 2002</u>; <u>Sarmiento-Rojas,</u> <u>2019</u>). During this interval, regional relative sea-level changes and paleoceanographic events primarily controlled the sedimentation in the basin (<u>Villamil and Pindell, 1998</u>). In the Maastrichtian–Early Paleocene interval, the final accretion of the Western Cordillera caused an initial inversion of inherited Mesozoic normal faults and structuration of Campanian strata

(<u>Cooper et al., 1995</u>; <u>Cooney and Lorente, 2009</u>). At the same time, uplifting of the Central Cordillera caused a change in the patterns of sedimentation transitioning from the marine Villeta Formation to the continental Rumiyaco Formation. Moreover, the convergence between the Nazca and South America plates intensified during the Early Paleocene–Middle Eocene, generating the inversion of extensional normal faults, and the development of thrust and back- thrust structures, erosion and deposition of the conglomerates of the Pepino Formation (<u>Casero et</u> <u>al., 1997</u>; <u>Córdoba et al., 1997</u>). During the Middle Eocene-Middle Miocene interval, the uplift of the Central Cordillera continued (<u>Cooper et al., 1995</u>) and induced flexural subsidence of the lithosphere, allowing the deposition of the Orito-Belén and Ospina formations. Ultimately, the tectonic inversion of Pre-Cretaceous, Cretaceous, and Paleogene depocenters is associated with the uplifting of the Eastern Cordillera and the development of the sub-Andean foreland basins (<u>Coletta et al., 1990</u>; <u>Cooper et al., 1995</u>; <u>Casero et al., 1997</u>).

## 3. Materials and methods

### 3.1. Locality

The 67 specimens of ammonoids and other macrofossils here studied (Figs 3–6) were collected *in-situ* during a field trip between January 7 and January 20, 2022, in a stratigraphic section cropping out along the Mocoa–San Francisco Road in the Putumayo Department, Colombia (Figs 1 and 2). This study is part of the AGESS project: “Astrocronología e Inversión Geoquímica aplicada a la identificación de “sweet spots” en rocas generadoras de clase mundial: Formaciones La Luna y Villeta, cuencas Putumayo y Valle Medio del Magdalena, Colombia”, funded by the Agencia Nacional de Hidrocarburos (ANH) and the Ministry of Science, Technology and Innovation of Colombia (Minciencias) by the grant No. 877-2021, and carried out by the Research Group Tectonics and Environmental Geology at the ‘Escuela de Ciencias Aplicadas e Ingeniería’, EAFIT University, Medellín, Colombia.

### 3.2. Materials

The studied material is held at the ‘Escuela de Ciencias Aplicadas e Ingeniería’ EAFIT University, Medellín, Colombia, with acronyms and catalogue numbers AGESS. The morphological study of 55 ammonoid samples in the area of study allowed us to recognize fourteen morphotypes and at least nine genera. The exact stratigraphic position of the samples is indicated in the stratigraphic column of the study area (Fig. 2, Suppl. Fig. S1).

For the anatomical description of the ammonoid specimens, we followed the works by <u>Klug et al. (2015)</u> and <u>Jain (2017)</u>, with emphasis in the recognition of features such as: type of umbilicum, type and direction of ribs, and type of ornamentation. The preservation of most ammonoid specimens here studied is as laterally-compressed shells, which in some cases prevents a detailed discussion at the specific or even generic levels. However, features such as delicate ornamentations, the diameter of the shells, and the umbilical dimensions, are well-preserved and allow a confident taxonomical assignation whenever possible. Based on the overall anatomical characters discernible in the studied specimens, we compared them with ammonoids known from other basins worldwide, including some from the Oriente Basin, Ecuador (e.g., <u>Tschopp, 1953</u>; <u>Bulot et al., 2005</u>), and the Upper Magdalena Valley Basin (UMVB) (e.g., <u>Etayo-</u> <u>Serna and Carrillo, 1996</u>), and their estimated stratigraphic ranges follow the works of <u>Wright et al.,</u> <u>1996</u>, <u>Villamil (1998)</u>, and <u>Kennedy and Klinger (2011)</u>.

### 3.3. Methods

Most of the macrofossils studied, particularly those from the Villeta Formation, are preserved in mudstones as semi-flattened impressions (Figs 3–5). No mechanical preparation was required. For macrophotography, the specimens were coated with sublimated ammonium chloride before photography to enhance relief of anatomical features and ornamentation. Specimens were imaged using a digital camera Canon EOS rebel T6.

## 4. Systematic Palaeontology

Phyllum: Mollusca <u>Linnaeus, 1758</u>

Class: Cephalopoda <u>Cuvier, 1797</u>

Order: Ammonitida <u>Hyatt, 1889</u>

Superfamily: Hoplitoidea <u>Douvillé, 1890</u>

Family: Schloenbachiidae <u>Parona and Bonarelli, 1896</u>

Genus: ***Schloenbachia*** <u>Neumayr, 1875</u>

? *Schloenbachia* sp.

Fig. 5A, P

*Material*. Two specimens, one with a maximum diameter of 70 mm (AGESS-P-0155-01) (Fig. 5A), and a smaller one with a diameter of 40 mm (AGESS-P-0155-12) (Fig. 5P).

*Description and remarks*. Involute coiling, with the umbilicus comprising about 25% of the diameter, less conspicuous in the small specimen. Fine ribbing and ventrolateral clavi are shared by both specimens; umbilical bullate tubercle is conspicuous in the large example. No keel observed.

*Remarks.* Colombian specimens are somehow comparable in size and tuberculation to involute less ornamented and compressed forms of *Schloenbachia* figured in <u>Kennedy (2013)</u>, but the ribbing is finer and without flank tubercles in the Colombian examples. *Mrhiliceras lapparenti* from Madagascar (<u>Wright et al., 1996</u>, fig. 117, 2c, d) shows a similar fine ribbing but bears a less marked umbilical tuberculation.

Family: Engonoceratidae <u>Hyatt, 1900</u>

Genus: ***Engonoceras*** Neumayr and Uhlig, 1881

cf. *Engonoceras* sp.

Fig. 5C, D, H–J, M

*Material*. Six specimens: AGESS-P-0155-03, -08, -11, -14, -15 and -21.

*Description*. Six fragmentary specimens each corresponding approximately to one third of a whorl. All small sized, except for one of them (AGESS-P-0155-03, Fig. 5C). Common

characters are high, slightly conical umbilical tubercles, and wide flexuous ribs with wider interspaces; primary ribs branch from the umbilical tubercle. In the smaller specimens (Fig. 5D, H–J, M) all ribs are weak or absent lower at the mid-flank, thicken towards the ventrolateral area where they show a claviform tubercle. The largest specimen figured does not show a ventrolateral clavi, the ventral area looks unornamented (despite its lateral compression) and the ribs are less flexuous than in the smaller ones (Fig. 5C).

*Remarks*. The ribbing type, coiling, and tuberculation, can be compared to *Engonoceras*- like ornamentation. However, the lack of clear ventral areas and conspicuous suture lines make difficult a better determination. If compared to *Engonoceras* forms from the upper Albian to lower Cenomanian of Texas (e. g., <u>Kennedy et al., 1998b</u>; <u>Kennedy et al., 2005</u>), the Colombian specimens share a robust ribbing but at smaller size.

Superfamily: Acanthoceratoidea de <u>Grossouvre, 1894</u>

Family: Brancoceratidae <u>Spath, 1934</u>

Subfamilia: Mojsisovicziinae <u>Hyatt, 1903</u>

Genus: ***Oxytropidoceras*** <u>Stieler, 1920</u>

Subgenus: Oxytropidoceras (Venezoliceras) <u>Spath, 1925</u>

Oxytropidoceras (Venezoliceras*) **sp***.

Fig. 3A–L, N–R

*Material.* Sixteen specimens: AGESS-P-0156-06, -07, -10, -12, -13, -17, -18, and -20, AGESS-P-0166-05, -09, -10, and -16, AGESS-P-0169-03, -04, and -06.

*Description.* Involute coiling and substantial whorl height increment is observable in the most complete specimen (Fig. 3E). Ribbing is well marked, with a density of about 25 ribs per half a whorl. Ribs are flexuous, bifurcate from a conspicuous umbilical bullate tubercle (e.g., Fig. 3I), get broader in the upper flank, and bend adorally in the ventrolateral shoulder where they are elevated and show the gradual acquisition of a strong ventrolateral claviform tubercle (e.g., Fig. 3C). Keel strong and elevated.

*Remarks*. Specimens AGESS-P-0166-09, -10, AGESS-P-0156-06, -10, -13, -18 and -20, show well-preserved flanks from the umbilical area to the keel; specimens AGESS-P-0156-07, - 12, -17, AGESS-P-0166-05 and -16, AGESS-P-0169-03, -04, -06 are partial flank remains that

preserve some taxonomically-informative characters. Coiling, ribbing pattern and the gradual acquisition of the ventrolateral tubercle make these specimens well identifiable as to be included in the subgenus *Venezoliceras*. The partial specimens are comparable to the best-preserved ones: ribbing pattern, keel, and specially the growth of the ventrolateral tubercle are characters observable among the specimens.

The well-marked ventrolateral tubercle of some specimens (Fig. 3C, I, K) and the ribbing pattern make the Colombian specimens comparable to *Oxytropidoceras* (*Venezoliceras*) *karsteni*, a species common in the upper Albian of Venezuela (<u>Renz, 1982</u>, pl. 17, figs. 2, 3).

Subgenus: Oxytropidoceras (Laraiceras) <u>Renz, 1968</u>

Oxytropidoceras (Laraiceras*) **sp***.

Fig. 3S, T

*Material.* One specimen: AGESS-P-0156-01, an almost complete one with flanks well preserved.

*Description*. Coiling and high keel similar to those of *O*. (*Venezoliceras*) sp. described above. Dense ribbing in the first half whorl preserved, with flexuous ribs bifurcating from a subtle bullate umbilical tubercle, being broader towards keel and showing a conspicuous tuberculate shape at the ventrolateral shoulder. This ribbing changes progressively towards the second half whorl becoming broader, with alternating long and short as well as bifurcating ribs near the mid flank, where a lateral tubercle appears; umbilical tubercle sharper and ventrolateral clavate tubercle strong specially in the last quarter preserved.

*Remarks*. The subgenus *Laraiceras* shows more spaced ribs and mainly alternating ones from the inner whorls. The specimen here described is placed in this subgenus due to the clearly developed trituberculate stage. Ribbing pattern at early growth stages with bifurcated ribs resembling *Venezoliceras* suggest that this specimen may be an early form of the subgenus.

cf. *Oxytropidoceras* sp.

Fig. 3M

*Material*. One specimen: AGESS-P-0159-08.

*Description.* Small specimen with a maximum 18 mm of diameter preserved, involute coiling, with an umbilicus comprising the 20% of the diameter. Keel is high, and the ventral area looks fastigiate on the last half a whorl, with a row of progressively sharper and claviform tubercles developed on the ventrolateral margin. Smooth flanks evolving to a very weak and spaced ribs on the last half a whorl.

*Remarks*. The well-formed ventrolateral tubercle, the high keel and the coiling make this Colombian specimen comparable to species of *Oxytropidoceras* (e. g., subgenus *Venezoliceras*), but in these, the tuberculated stage occurs later in the ontogeny together with a well marked ribbing. The development of sharp tubercles, the change of shape of the ventral area and the mainly smooth flanks at a very small size indicate the fast acquisition of an overall changing ornamentation at early ontogeny, and thus this specimen could be considered a dwarf form.

Other features such as the presence of crowded suture lines towards the end of the conch would help to determine if this is a dwarf offshoot of *Oxytropidoceras*.

Subfamily: Mortoniceratinae <u>Douvillé, 1912</u>

Genus: ***Mortonicera****s* <u>Meek, 1876</u>

Subgenus: Mortoniceras (Mortoniceras) <u>Meek, 1876</u>

*Discussion*. In a recent paper on upper Albian ammonites from North-East Texas, <u>Gale</u> <u>and Kennedy (2020)</u> discuss the differences between two lineages of mortoniceratines, one with the characters of the *vespertinum* species, and the other composed of cosmopolitan forms allied to the *inflatum-rostratum* group, named *Mortoniceras* by previous authors. Accordingly, they propose the use of the genus *Mortoniceras* for the *vespertinum* group, and to recover the genus *Pervinquieria* for the other one. Given the few mortoniceratin specimens at hand from the stratigraphic section here studied, we follow the nomenclature from the Treatise (<u>Wright et al.,</u> <u>1996</u>) until new Colombian material becomes available for a more in-depth discussion of their generic placement.

Mortoniceras (Mortoniceras*) **cf.*** vespertinum

Fig. 4D

2020 *Mortoniceras vespertinum* (<u>Morton, 1834</u>): <u>Gale and Kennedy, 2020</u>, p. 30; plate XI. Fig. 7-9 (with synonymy).

*Material*. One specimen: AGESS-P-0166-04.

*Description*. Specimen with a maximum 32 mm of diameter preserved. Evolute coiling with umbilicus comprising about 40% of the diameter. More than the last half a whorl bearing sharp conical umbilical and broader-claviform ventrolateral tubercles gaining strength adorally. Ribs very weak, possibly due to preservation, but clearly not well marked on the flank; they look straight, with primaries and maybe secondaries at the end of the shell, but not branching is evident. The ventrolateral area seems to bear a division of the tubercle at the end of the preserved whorl. Keel is only conspicuous in the last quarter of the whorl.

*Remarks*. Despite its preservation, coiling, ribbing style and the tubercle shape and arrange are key features observed in innermost whorls of species of the *vespertinum* group. The lack of adult stages in the Colombian collection prevents additional determinations; however, it is quite comparable to the early growth stages at similar diameter observed in *vespertinum* specimens of North Texas (e.g., <u>Gale and Kennedy, 2020</u>, pl. XI, figs. 7–9).

Mortoniceras (Mortoniceras*) **sp***.

Fig. 4A–C

*Material*. Two specimens: AGESS-P-0168, near half a whorl of what seems a partial body chamber (Fig. 4A, B), and AGESS-P-0169-02, a fragmentary upper flank (Fig. 4C).

*Description*. Specimen AGESS-P-0168 shows near 14 wide ribs separated by similarly wide interspaces, with a shape suggesting an evolute coiling. Ribs slightly flexuous, mainly primaries extending from the umbilical wall to the ventrolateral area where they are broader but do not reach the keel; some secondaries arise from the lower flank. Umbilical bullate tubercle high; mid-flank tubercle low, and ventrolateral tubercle bullate, broader near the keel. Ribs show conspicuous spiral ridges. The other specimen, AGESS-P-0169-02, fits this description.

*Remarks*. Spiral ridges are a main feature of the genus *Elobiceras*, but it is observed also in other mortoniceratines (e. g. subgenus *Mortoniceras* and *Deiradoceras*). The presence of a mid-flank tubercle clearly separates these specimens from *Elobiceras* and from bituberculated

species of *Mortoniceras*. The trituberculation ribbing and coiling make the Colombian specimens be compared to forms of the *inflatum* group. Coarse ribbed trituberculated forms have been described in Texas (e.g., <u>Gale and Kennedy, 2020</u>) or in Europe (e.g. <u>López-Horgue et al., 1999</u>), some of them with spiral ridges, but the fragmentary colombian sample prevents further discussion.

Genus: ***Algericeras*** <u>Spath, 1925</u>

Algericeras ***sp*.**

Fig. 5R–U

*Material*. Four specimens: Three complete small specimens with a similar diameter of about 15 mm, AGESS-P-0155-17, -18, and -19, and one fragment comprising a third of a whorl, AGESS-P-0155-20.

*Description*. Umbilicus comprise a bit less than 30% of the diameter. Considering their small size, the ribbing is well developed in the last whorl showing similar characteristics. Ribbing is dense with about 30 flexuous ribs in the last half a whorl. Ribs are sharp and narrow, with narrower interspaces; primaries branch into two from a slightly bullate umbilical tubercle, being projected adorally at the ventral area from the upper third of the flank, where they are the widest. Secondary ribs arise from at lower third of the flank. Keel is not well preserved due to compression, but it is conspicuous at different parts on the ventral end.

*Remarks*. Despite the moderate degree of preservation of the Colombian specimens, their coiling, umbilical breadth, ribbing, and keel, make them comparable to *Algericeras*. This genus is composed of small, dwarf examples that show a developed ribbing at small diameters such as those of the samples here studied. In fact, the Colombian specimens are comparable to the type specimens of *Algericeras bogharensis* <u>Coquand, 1879</u>, from the type locality in Northern Algeria, and other related from Tunisia (e.g., <u>Kennedy, 2020</u>, pl. 12, figs. 1-15, and pl. 13, figs. 6–8).

Subfamily: Brancoceratinae <u>Spath, 1934</u> Genus: ***Hysteroceras*** Hyatt, 1900

*Hysteroceras* sp. 1

Fig. 4I, J

*Material*. Two specimens: AGESS-P-0166-12 and AGESS-P-0166-17.

*Description*. Both specimens small sized, quite evolute, with an umbilicus around one third of a 20 mm in diameter. Sharp and dense ribbing, with near 20 ribs in the last half whorl. Ribs are flexuous, look broader than interspaces; they extend from the umbilical area showing a sharp umbilical tubercle from which rib bifurcation can be observed, cross the flank and become broader at the ventrolateral area sharply bending towards a low keel. On the last half a whorl some solitary ribs are intercalated between branching ones.

*Remarks*. The size, the coiling, the apparently quite conservative ribbing, and the sharp umbilical tubercle, are features shared by many species of *Hysteroceras*. Rib density, their flexuous shape, and the presence of a low keel in the late stage, are features that can be seen in species like *H. carinatum* from Venezuela (e.g. <u>Renz, 1982</u>, pl. 11, figs. 9–11), but Colombian specimens seem to be slightly less evolute and more comparable to Tethyan forms (e.g. <u>López-</u> <u>Horgue et al., 2009</u>, fig. 6 D–F).

*Hysteroceras* sp. 2

Fig. 4H, M

*Material*. Two specimens: AGESS-P-0156-04 and AGESS-P-0167-01.

*Description*. Small sized, with a diameter of 41 and 28 mm, respectively. Evolute coiling, with the umbilicus comprising the 44% of the diameter in both specimens. Ribs well developed from the innermost observable parts, quite conservative throughout the shell, consisting on slightly flexuous sharp ribs, bearing a sharp umbilical tubercle from which ribs branch; ribbing looks not branched at the end of the shell in specimen 0156-04 (fig. 4 H), with a conspicuous elevated venter without keel. Ventrolateral terminations do not show a conspicuous tubercle at any growth stage.

*Remarks*. Both specimens share similar ribbing and coiling, thus they are included in the same taxonomic determination. Smaller one is interpreted as a juvenile with always branched ribbing. The ribbing pattern and tuberculation, as well as size, are compatible with those of *Hysteroceras* species. Specimens are somehow comparable to some species without keel at the

end of the conch occurring in the upper Albian of Venezuela (e.g., <u>Renz, 1982</u>, pl. 11, fig. 12, and pl. 12, fig. 3).

cf. *Hysteroceras* sp.

Fig. 4G, K, L

*Material*. Three specimens: AGESS-P-0156-19, -21, and AGESS-P-0158-06.

*Description*. Small conchs with diameters from 20 to 35 mm and wide umbilicus around 40% of the diameter, with a ribbing pattern similar to described *Hysteroceras*; keel is only partially visible at the early growth stage in one of them (AGESS-P-0156-21). Conservation avoids better determinations.

Family: Forbesiceratidae <u>Wright, 1952</u>

Genus: ***Forbesiceras*** <u>Kossmat, 1897</u>

cf. *Forbesiceras* sp.

Fig. 5F, G, K, L, N, O

*Material*. Six specimens: Three fragmentary specimens partially showing the upper flank and the ventral area; AGESS-P-0155-04, -09, and -22, and three fragmentary specimens showing the coiling of the last preserved whorl; AGESS-P-0155-05, -07, and -10.

*Description and remarks*. Despite the preservation, the ribs broadening towards the venter and mainly developed in the upper flank, together with the ventrolateral clavi linked by a subtle transversal rib with delicate parallel striations (see Fig. 5K), make these Colombian specimens comparable to specimens of the genus *Forbesiceras* (e.g., the Algerian *F. largilliertianum* in <u>Kennedy, 2020</u>, and the Tunisian *F. chevillei* in <u>Kennedy and Gale, 2015</u>).

Family: Acanthoceratidae de <u>Grossouvre, 1894</u>

Subfamily: Mantelliceratinae <u>Hyatt, 1903</u>

Genus: ***Graysonites*** <u>Young, 1958</u>

***Graysonites*** sp.

Fig. 5B, E, Q

*Material*. Three specimens: AGESS-P-0155-02 (one whorl preserved), AGESS-P-0155- 06, and AGESS-P-0155-24 (both fragmentary, approximately half a whorl); all of very similar size.

*Description*. Involute coiling with high whorl expansion; in the complete specimen, the umbilicus comprises less than 25% of a 25 mm diameter. Flexuous ribbing is better preserved in specimen 0155-06 with a conspicuous umbilical bullate tubercle that gives rise to primary ribs crossing the flank and showing a well-marked inner and clavate outer ventrolateral tubercles.

Secondary ribs between primaries do not reach the umbilical area and the interspaces are of similar width. There is no clear rib branching except for specimen 0155-24.

*Remarks*. Despite the ventral area is not observed due to conservation, coiling, bullate umbilical tubercle and the row of two ventrolateral tubercles are diagnostic features of the genus *Graysonites*. Specimens are comparable to similar size examples from Texas (<u>Kennedy et al.,</u> <u>2005</u>, figs 27T, 28P, 29A, B), Tunisia (<u>Kennedy and Gale, 2015</u>, pl. IX, figs. 4, 8) and Algeria (<u>Kennedy, 2020</u>, pl. 22, fig. 7). Colombian specimens show flexuous ribs as in the Algerian *Graysonites elegans* <u>Kennedy, 2020</u>.

Superfamily: Turrilitoidea <u>Gill, 1871</u>

Family: Hamitidae <u>Gill, 1871</u>

Genus: ***Hamites*** <u>Parkinson, 1811</u>

***Hamites*** sp.

Fig. 4N–Q

*Material*. Four specimens: AGESS-P-0156-02, and AGESS-P-0166-01, -02, -03.

*Description*. Coiling in one plane is observable in specimen AGESS-P-0156-02 (Fig. 4Q), which is a juvenile with open plane spiral coiling changing to a more open shaft adorally; rectiradiate ribbing weak. The rest of the specimens are partial shafts with a very open curvature in two of them (Fig. 4N, P), all showing rectiradiate ribbing, with approximately equal width in ribs and interspaces. No conspicuous tubercles.

*Remarks*. Fragmentary hamitids are difficult to ascribe to a given species, even to a genus, but the studied specimens compare well to open coiled *Hamites* species on the basis of their

ribbing and open coiling. The juvenile specimen shows a similar coiling as *Idiohamites* at the same growth stage, but the lack of tubercles prevents its assignation to this genus.

## 5. Results

### 5.1. Age of the Ammonoid Bearing Intervals

In general terms, we distinguish two clearly separated ammonoid-bearing intervals in the Putumayo section with differentiated faunal composition. The lower interval extends from the meters 153.5 to 170 (Fig. 2, Suppl. Fig. S1), and is dominated by elements of the family Brancoceratidae, with minoritary hamitids. Six ammonoid-bearing levels have been identified in this interval (Fig. 2) showing the following faunal succession from the lowermost one upwards: 1) *Hysteroceras* sp.; 2) cf. *Oxytropidoceras* sp.; 3) *Oxytropidoceras* (*Venezoliceras*) sp., *Hysteroceras* sp. 1, *Mortoniceras* (*Mortoniceras*) cf. *vespertinum*, and *Hamites* sp.; 4) *Hysteroceras* sp. 2.; 5) *Mortoniceras* (*Mortoniceras*) sp.; and 6) *Oxytropidoceras* (*Venezoliceras*) sp., *Oxytropidoceras* (*Laraiceras*) sp., *Hysteroceras* sp. 2, cf. *Hysteroceras* sp., *Mortoniceras* (*Mortoniceras*) sp. and *Hamites* sp. (Figs 3, 4). As a whole, this interval is of early late Albian age, based on the presence of the cosmopolitan genus *Hysteroceras* and *Hamites*, and the subgenus *Venezoliceras*, *Laraiceras* and *Mortoniceras* (Fig. 7).

There is a nearly 100 meter-thick gap in the ammonoid succession mainly due to poor outcrop exposition, therefore the upper ammonoid interval in the Putumayo section is the single level 7) at stratigraphic meter 271 (Fig. 2). This level bears the following association: ?

*Schloenbachia*, cf. *Engonoceras* sp., cf. *Forbesiceras* sp., *Algericeras* sp. and *Graysonites* sp. (Fig. 5). This assemblage characterizes the lowermost Cenomanian. Other fossils, such as the fish remains (e.g., scales) and the plant remains are too fragmentary or non-diagnostic as to elaborate on their detailed systematic affinities. The decapod crustacean remain (Fig. 6L) seems to correspond to a callianassioid axiidean ghost shrimp. While a more precise systematic placement is not possible at this time, it is worth mentioning that this is the first report of crustacean macrofossils from the Putumayo Basin.

## 6. Discussion

### 6.1. The upper Albian ammonoid association

According to the described specimens, the upper Albian ammonoid associations show typical elements from Venezuela (*Venezoliceras*, *Laraiceras;* e.g., <u>Renz, 1982</u>) and Texas (*Mortoniceras* (*M.*) cf. *vespertinum;* e.g., <u>Gale and Kennedy, 2020</u>) together with cosmopolitan *Hysteroceras* and *Hamites*, characteristics of shallow marine platforms of the Tethyan province extending from South America to Western Europe, North Africa, Madagascar, India, and Iran, during early Late Albian times (e.g. <u>Owen and Mutterlose, 2006</u>). *Venezoliceras* has also been described from Western Europe (e.g., <u>López-Horgue et al., 2009</u>), Peru, Texas, Mexico, Africa and Madagascar (<u>Wright et al., 1996</u>). According to this Tethyan distribution, we follow here the upper Albian ammonoid zonal scheme proposed for the Western Tethyan area (in Northern Spain, <u>López-</u> <u>Horgue et al., 1999</u>; <u>López-Horgue et al., 2009</u>; and in Suriname, <u>Owen and Mutterlose, 2006</u>); this scheme is mainly based on the ranges of brancoceratid cosmopolitan elements of widespread occurrence and relative abundance, of the genus *Hysteroceras* and *Mortoniceras*.

The Lower Cretaceous ammonite working group (the Kilian Group) proposed an upper Albian biozonation based on the phyletic evolution of Mortoniceratines (<u>Reboulet et al., 2011</u>), but some index species are very scarce and need revision (<u>López-Horgue and Owen, 2022</u>). The use of *Hysteroceras* and other elements is especially interesting for the lower part of the upper Albian (the former *M. inflatum* Zone of <u>Spath, 1941</u>). Besides, the transition to the uppermost Albian (to the former *Stoliczkaia dispar* Zone) is also a matter of debate. The occurrence of *M.* (*M.*) cf. *vespertinum* in Putumayo and the Tethyan elements in Texas let us to consider an attempt of correlation with the biozonation in the Texas area, and to compare the Western Tethyan biozonation used here to the tentative biozonal correlation of North-East Texas and Western Europe proposed by <u>Gale and Kennedy (2020)</u> (Fig. 7).

In Texas, the first Zone of *Adkinsites bravoensis* contains *Hysteroceras varicosum* and *Deiradoceras* spp., which are also a common faunal element in the *Hysteroceras orbignyi* and *H. binum* Subzones in the Western Tethys (e.g., <u>López-Horgue et al., 2009</u>). The occurrence in Texas of the inoceramid *Actinoceramus sulcatus* atop of the Kiamichi Formation (forma C, synonym of *A. sulcatus biometricus*; <u>Kennedy et al., 1999</u>), i. e., the top of *A. bravoensis* Zone (see fig. 4, <u>Gale</u>

<u>and Kennedy, 2020</u>), is correlatable with the transition to the *H. binum* Zone in Europe (e. g., see fig. 1-5 of <u>Gale and Owen, 2010</u>). In Putumayo, occurrences in the first two levels are not enough to give a precise biozone, and so it is given by the stratigraphic order underneath level 3 (Fig. 2), as discussed below.

The *Elobiceras* (*Craginites*) *serratescens* Zone in Texas contains *Hysteroceras* cf. *varicosum* and *H.* cf. *binum*; the latter is the marker of its homonym Subzone in the Tethys (e.g., <u>Owen and Mutterlose, 2006</u>). *Craginites* sp. is a component of the *H. choffati* Subzone in Northern Spain (<u>López-Horgue et al., 2009</u>). *Mortoniceras* (*M.*) *vespertinum* occurs in the upper part of the *Eopachydiscus marcianus* Zone of Texas above the *E.* (*C.*) *serratescens* Zone. In Putumayo *M.* (*M.*) cf. *vespertinum* occurs together with *Oxytropidoceras* (*Venezoliceras*) sp. and *Hysteroceras* sp. 1 in the level 3, above the levels 1 and 2 with only cf. *Hysteroceras* sp. and cf.

*Oxytropidoceras* sp. Accordingly, levels 1 and 2 in Putumayo may lie below sediments of the transition *H. choffati* to *M.* (*M.*) *inflatum* Zones (level 3, Fig. 2), corresponding to the *Hysteroceras varicosum* Zone without further precision (Figs 2 and 7). *Pervinquieria* (*P.*) *equidistans* Zone in Texas contains mortoniceratines with a clear mid-flank tubercle, which are a distinctive element of the *M.* (*M.*) *inflatum* Zone in Europe (e.g., <u>Owen, 2012</u>). *Mortoniceras* (*M.*) *sp.* with a conspicuous mid-flank tubercle occur in the levels 5 and 6 of Putumayo together with cf. *Hysteroceras* sp. and *H.* sp. 2 in the level 6, indicating probably the lower *Mortoniceras* (*M.*) *inflatum* Zone (sensu <u>Wiedmann and Owen (2001)</u> and <u>Owen (2012)</u>) at least between m. 166 and 170 (levels 5 and 6; Figs 2 and 7). Abundant *Oxytropidoceras* (*Venezoliceras*) sp. and *O.* (*Laraiceras*) sp. occur also in the level 6 of Putumayo, supporting a topmost record of this genus and subgenus in Colombia.

### 6.2. The lower Cenomanian ammonoid association

The Putumayo level 7 (Fig. 2) contains quite a varied association of typical mainly Tethyan Cenomanian forms. *Graysonites* is a genus whose type species comes from Texas, but shows a widespread occurrence during the Early Cenomanian (Texas, California, Brazil, Tunisia, Algeria, Japan, Spain) (e. g. <u>Wright et al., 1996</u>; <u>Kennedy, 2020</u>). The Putumayo *Graysonites* specimens are mainly comparable to some Algerian forms. *Algericeras* sp. from Putumayo is a form comparable also to the Algerian ones (<u>Kennedy, 2020</u>); the genus characterizes the basal Cenomanian also in Tunisia, Madagascar, and Mexico. *Schloenbachia* is a highly plastic form

that characterizes the lower Cenomanian in the European province; the Putumayo record is not suitable for further discussion. The rest of the association is composed of cf. *Forbesiceras* sp. and cf. *Engonoceras* sp., a longer ranging forms; the former ranges from the lowermost Cenomanian (*N. carcitanense* Subzone of the *M. mantelli* Zone) to the uppermost Cenomanian *M. geslinianum* Zone in Europe (<u>Kennedy and Klinger, 2008</u>), and the latter is a common element in the lower Albian to lower Cenomanian associations in Texas (e. g., <u>Kennedy et al., 1998a</u>; <u>Kennedy et al.,</u> <u>1998b</u>; <u>Kennedy et al., 2005</u>), and shows a wider occurrence (Europe, Algeria, Tunisia, Mexico and Colombia).

### 6.3. The Albian–Cenomanian boundary in northwestern South America

The Albian–Cenomanian boundary at 100.5 Ma coincides not only with the boundary of the foraminifera zones *Parathalmanninella appenninica* and *Thalmanninella globotruncanoides*, but also with the first occurrence of the calcareous nannoplankton *Corollithion kennedyi* (<u>Gale et al.,</u> <u>2020</u>). In northwestern South America, the *Parathalmanninella appenninica* biozone has been recognized in rocks of the Querecual Formation consistent with the presence of upper Albian rocks in Eastern Venezuela (<u>Crespo de Cabrera et al., 1999</u>). The Early Cenomanian *Thalmanninella globotruncanoides* biozone, on the other hand, has not been identified in northwestern South America, hampering the unequivocal assignation of rock formations to this age stage.

In northwestern South America, specifically in Venezuela and Colombia, <u>Renz (1982)</u> and <u>Villamil (1998)</u> recognized that the lowermost part of the Cenomanian is hard to recognize based on ammonites. This interpretation is apparently consistent with seismic, stratigraphic, and paleontological evidences that point out to the presence of an unconformity between the Albian and Cenomanian in several basins of the northern Andes (<u>Jaillard et al., 2000</u>; <u>Jaimes and de Freitas, 2006</u>; <u>Etayo-Serna, 2019</u>). Ammonites and foraminifera in the Upper Magdalena Valley Basin of Colombia, however, indicate that the Albian–Cenomanian boundary is present within the lower part of the “Shale y bancos con *Costagyra*” informal unit (<u>Etayo-Serna and Carrillo, 1996</u>; <u>Leon</u> <u>Rodriguez, 2002</u>). Similarly, ammonoid and bivalve faunas in Peru show that the Albian– Cenomanian boundary is identifiable in the central and northern part of the country (<u>Benavides-</u> <u>Cáceres, 1956</u>; <u>Navarro-Ramirez et al., 2017</u>).

The ammonoid-dominated macrofossil fauna from the Putumayo Basin here studied span the late Albian and the early Cenomanian, and serve as a point of reference to compare neighboring basins (e.g., Marañon in Perú, Oriente in Ecuador, Upper Magdalena Valley in Colombia) (Fig. 1), and thus the generation of more comprehensive biostratigraphic correlations for northwestern South America. Namely, the occurrence of the ammonoid genera *Oxytropidoceras*, *Mortoniceras*, and *Schloenbachia*, suggest interconnection during the mid-Cretaceous of the Putumayo Basin with the Upper Magdalena Valley in Colombia and the Oriente Basin in Ecuador (<u>Tschopp, 1953</u>; <u>Etayo-Serna and Carrillo, 1996</u>; <u>Bulot et al., 2005</u>).

## 7. Conclusions

- We report 67 new *in-situ* occurrences of upper Albian–lower Cenomanian (∼105–96 Ma) ammonoids, bivalves, decapod crustaceans, and other macrofossils, from the Villeta Formation in Putumayo, Colombia, a time interval and geographic area poorly known in northwestern South America due to poor exposition of rocks and apparent stratigraphic hiatuses between the upper Albian and the lower Cenomanian.
- We recognized two main stratigraphic intervals containing distinctive ammonoid associations. The lowermost interval includes six levels bearing ammonoids such as *Hysteroceras* spp., *Oxytropidoceras* sp., *O.* (*Venezoliceras*) sp., *O.* (*Laraiceras*) sp., *Mortoniceras* (*Mortoniceras*) cf. *vespertinum*, *M.* (*Mortoniceras*) sp., and *Hamites* sp., indicative of the upper Albian, whereas the uppermost interval includes one ammonoid- bearing level with*Schloenbachia*, cf. *Engonoceras* sp., cf. *Forbesiceras* sp., *Algericeras* sp. and *Graysonites* sp., indicative of the lowermost Cenomanian.
- These mid-Cretaceous ammonoid assemblages fill a spatial and temporal gap in our understanding of the Putumayo Basin and its fossil record, and will facilitate future biostratigraphic correlations with neighbouring basins in northwestern South America (e.g., Upper Magdalena Valley and Llanos basins in Colombia, Cuenca Oriente in Ecuador, Marañón Basin in Perú), as well as to other mid-Cretaceous basins worldwide.

## Acknowledgements

This study was supported by the “Convocatoria para la Financiación de Proyectos de Investigación en Geociencias para el Sector de Hidrocarburos, número: 877”, by the Agencia Nacional de Hidrocarburos (ANH) and the Ministerio de Ciencia, Tecnología e Innovación de Colombia (Minciencias), grant No. 877-2021, as part of the AGESS project: “Astrocronología e Inversión Geoquímica aplicada a la identificación de “sweet spots” en rocas generadoras de clase mundial: Formaciones La Luna y Villeta, cuencas Putumayo y Valle Medio del Magdalena, Colombia”, carried out by the Research Group Tectonics and Environmental Geology at the ‘Escuela de Ciencias Aplicadas e Ingeniería’, EAFIT University, Medellín, Colombia. Special thanks to Andrés Cárdenas (EAFIT) for early discussions and suggestions, to Laura Redondo and Cristian Valencia (EAFIT), and Camilo Rengifo for their assistance in the field. We thank XX and XX reviewers for their suggestions and feedback on an earlier version of this manuscript.

## Author Contribution

**Javier Luque**: Conceptualization, Investigation, Writing – Original Draft preparation, Visualization, Supervision, Funding acquisition. **Jonatan Bustos**: Conceptualization, Investigation – Fieldwork, Writing – Original Draft preparation, Visualization, Supervision. **Alejandro Beltrán-Triviño**: Writing – Original draft preparation, Visualization, Supervision, Funding acquisition, Project Administration. **Angie Rodriguez**: Investigation – Fieldwork, Project Administration. **Alexandra Delgado**: Investigation – Fieldwork. **Johan M. Sanchez**: Investigation – Fieldwork. **Julián Caraballo**: Investigation, Writing – Original Draft preparation. **Manuel Paez-Reyes**: Investigation – Fieldwork, Writing – Review & Editing, Funding acquisition. **Mikel A. López-Horgue**: Conceptualization, Investigation, Writing – Original Draft preparation, Visualization. All authors have read and approved the submission of the manuscript.

## Data and materials availability

All data are present in the paper and as Supplementary Materials.

## Tables

Table 1. List of mid-Cretaceous macrofossils (ammonoids, bivalves, crustacean remains, fish remains, plant remains) here studied from the middle–upper Aptian to lower Cenomanian Caballos and Villeta formations in the Mocoa–San Francisco section, Putumayo region, Colombia, northern South America). Stratigraphic heights as in Fig. 2 and Supplementary Appendix S1.

## Supplementary Figure S1

**Supplementary Figure S1.** Detailed lithostratigraphic column of the mid-Cretaceous (Albian–Cenomanian) section studied in the Mocoa–San Francisco road, Department of Putumayo, Colombia, northern South America.

## References

1. Benavides-Cáceres, V.E., 1956. Cretaceous system in northern Peru. Bulletin of the American Museum of Natural History 108, 1–212.

2. Bulot, L.G., Kennedy, W.J., Jaillard, E., Robert, E., 2005. Late Middle-early Late Albian ammonites from Ecuador. Cretaceous Research 26, 450–459.

3. Casero, P., Salel, J.F., Rossato, A., 1997. Multidisciplinary correlative evidences for polyphase geological evolution of the foothills of the Cordillera Oriental (Colombia). Proceedings of the VI Simposio Bolivariano, Cartagena, Colombia, 100–118.

4. Coletta, B., Hebrard, F., Letouzey, J., Werner, P., Rudkiewicz, J.L., 1990. Tectonic style and crustal structure of the Eastern Cordillera (Colombia) from a balanced cross-section, in: Letouzey, J. (Ed.), Petroleum and tectonics in mobile belts. Editions Technip, Paris, pp. 81–100.

5. Cooney, P., Lorente, M., 2009. A structuring event of Campanian age in western Venezuela, interpreted from seismic and palaeontological data. Geological Society, London, Special Publications 328, 687–703.

6. Cooper, M., Addison, F., Alvarez, R., Coral, M., Graham, R., Hayward, A., Howe, S., Martinez, J., Naar, J., Peñas, R., Pulham, A.J., Taborda, A., 1995. Basin development and tectonic history of the Llanos Basin, Eastern Cordillera, and middle Magdalena Valley, Colombia. AAPG bulletin 79, 1421–1443.

7. Coquand, H., 1879. Études supplémentaires sur la paléontologie algérienne. Bulletin de l’Académie d’Hippone 15, 1–449.

8. Córdoba, F., Buchelli, F., Moros, J., Calderón, W., Guerrero, C., Kairuz, E.C., Magoon, L., 1997. Proyecto evaluación regional Cuenca del Putumayo—Definición de los sistemas petrolíferos, Ecopetrol, Internal Report, p. 140.

9. Crespo de Cabrera, S., Sliter, W.V., Jarvis, I., 1999. Integrated foraminiferal biostratigraphy and chemostratigraphy of the Querecual Formation (Cretaceous), eastern Venezuela. The Journal of Foraminiferal Research 29, 487–499.

10. Cucalón, I., Camacho, R., 1966. Compilación geológica de la cuenca del Putumayo, pp. 1–18.

11. Cuvier, G., 1797. Tableau élémentaire de l ’ histoire naturelle des animaux. Baudouin, imprimeur.

12. Douvillé, H., 1890. Sur la classification des Cératites de la Craie. Impr. Bigot frères.

13. Douvillé, H., 1912. Évolution et classification des Pulchelliidés. Protat Frères.

14. Etayo-Serna, F., 2019. Basin development and tectonic history of the Middle Magdalena Valley, Estudios geológicos y Paleontológicos sobre el Cretácico en la Región del Embalse del Río Sogamoso, Valle Medio del Magdalena. Servicio Geológico Colombiano, Bogotá, pp. xx-xx.

15. Etayo-Serna, F., Carrillo, G., 1996. Bioestratigrafía del Cretácico mediante Macrofósiles en la Sección El Ocal, Valle Superior del Magdalena, Colombia. Geología Colombiana 20, 81–92.

16. Gale, A., Mutterlose, J., Batenburg, S., Gradstein, F., Agterberg, F., Ogg, J., Petrizzo, M., 2020. The Cretaceous Period, Geologic time scale 2020. Elsevier, pp. 1023–1086.

17. Gale, A.S., Kennedy, W.J., 2020. Upper Albian ammonites from North-East Texas. Revue de Paleobiologie 39, 1–139.

18. Gale, A.S., Owen, H.G., 2010. Introduction to the Gault, in: Young, J.R., al., e. (Eds.), Fossils of the Gault Clay. The Palaeontological Association, London, pp. 1–15.

19. Gill, T., 1871. Arrangement of the families of mollusks prepared for the Smithsonian Institution (Vol. 227). Smithsonian Institution.

20. Gonçalves, F.T., Mora, C.A., Córdoba, F., Kairuz, E.C., Giraldo, B.N., 2002. Petroleum generation and migration in the Putumayo Basin, Colombia: insights from an organic geochemistry and basin modeling study in the foothills. Marine and petroleum geology 19, 711-725.

21. Grossouvre, A.D., 1894. Recherches sur la craie supérieure, 2, Paléontologie. Les ammonites de la craie supérieure. Mémoires du Service de la Carte géologique détaillée de la France.

22. Hyatt, A., 1889. Genesis of the Arietidae. Smithsonian Contributions to Knowledge 673, Washington, DC xi, 16.

23. Hyatt, A., 1900. Cephalopoda, in: Zittel, K.A.V. (Ed.), Textbook of Palaeontology. Eastman, London and New York, pp. 502–592.

24. Hyatt, A., 1903. Pseudoceratites of the Cretaceous (Vol. 44). US Government Printing Office.

25. Jaillard, E., Hérail, G., Monfret, T., Díaz-Martínez, E., Baby, P., Lavenu, A., Dumont, J.-F., Cordani, U., Milani, E., Campos, D., 2000. Tectonic evolution of the Andes of Ecuador, Peru, Bolivia and northern Chile, in: Cordani, U., Milani, E., Thomaz Filho, A., Campos, D. (Eds.), Tectonic Evolution of South America, Rio de Janeiro, Brazil, pp. 481–559.

26. Jaimes, E., de Freitas, M., 2006. An Albian–Cenomanian unconformity in the northern Andes: Evidence and tectonic significance. Journal of South American Earth Sciences 21, 466–492.

27. Jain, S., 2017. Cephalopoda, Fundamentals of Invertebrate Palaeontology. Springer India, pp. 31–102.

28. Kennedy, W.J., 2013. On variation in *Schloenbachia varians* (J. Sowerby, 1817) from the Lower Cenomanian of western Kazakhstan. Acta Geologica Polonica 63, 443–468.

29. Kennedy, W.J., 2020. Upper Albian, Cenomanian, and Upper Turonian ammonite faunas from the Fahdène Formation of Central Tunisia and correlatives in northern Algeria. Acta Geologica Polonica 70, 147–272.

30. Kennedy, W.J., Cobban, W.A., Gale, A.S., Hancock, J.M., Landman, N.H., 1998a. Ammonites from the Weno Limestone (Albian) in Northeast Texas. American Museum Novitates 3236, 46 p. 3236, 1–46.

31. Kennedy, W.J., Cobban, W.A., Hancock, J., Gale, A.S., 2005. Upper Albian and Lower Cenomanian ammonites from the Main Street Limestone, Grayson Marl and Del Rio Clay in northeast Texas. Cretaceous Research 26, 349–428.

32. Kennedy, W.J., Gale, A.S., 2015. Upper Albian and Cenomanian ammonites from Djebel-Mhrila, Central Tunisia. Revue de Paleobiologie 34, 235–361.

33. Kennedy, W.J., Gale, A.S., Hancock, J.M., Crampton, J.S., Cobban, W.A., 1999. Ammonites and inoceramid bivalves from close to the Middle-Upper Albian boundary around Fort Worth, Texas. Journal of Paleontology 73, 1101–1125.

34. Kennedy, W.J., Klinger, H.C., 2008. Cretaceous faunas from Zululand and Natal, South Africa. The ammonite family Forbesiceratidae Wright, 1952. African Natural History 4, 117–130.

35. Kennedy, W.J., Klinger, H.C., 2011. Cretaceous faunas from Zululand and Natal, South Africa. The ammonite genus Oxytropidoceras Stieler, 1920. African Natural History 7, 69-102.

36. Kennedy, W.J., Landman, N.H., Cobban, W.A., 1998b. Engonoceratid ammonites from the Glen Rose limestone, Walnut clay, Goodland limestone, and Comanche Peak limestone (Albian) in Texas. American Museum Novitates 3221, 1–40.

37. Kennedy, W.J., Wright, C.W., 1981. *Euhystrichoceras* and *Algericeras*, the last Mortoniceratine ammonites. Palaeontology 24, 417–435.

38. Klug, C., Korn, D., Landman, N.H., Tanabe, K., Baets, K.D., Naglik, C., 2015. Describing ammonoid conchs, in: Klug, C., Korn, D., De Baets, K., Kruta, I., Mapes, R. (Eds.), Ammonoid Paleobiology: from anatomy to ecology (Vol. 43). Springer, pp. 3–24.

39. Kossmat, F., 1897. Untersuchungen uber die Sudindische Kreideformation. Beitrage zur Palaontologie und Geologie Osterreich-Ungarns und des Orients 11, 1–46.

40. Leon Rodriguez, L., 2002. Estudio micropaleontológico y bioestratigráfico con base en foraminíferos planctónicos de las "Calizas del Tetuán, Grupo Villeta". Valle Superior del Magdalena. Universidad Nacional de Colombia, Bogotá, p. 279.

41. Linnaeus, C.v., 1758. Systema Naturae per Regna Tria Naturae, Secundum Classes, Ordines, Genera, Species, cum Characteribus, Differentiis, Synonymis, Locis. Laurentii Salvii, Holmiae.

42. López-Horgue, M.A., Owen, H.G., 2022. Mortoniceratinae (Cretaceous ammonitina) from the Basque-Cantabrian Basin (BCB, Western Pyrenees): a key to understanding Upper Albian biostratigraphy, in: Jagt, J.W.M.e.a. (Ed.), Abstract volume of the 11th International Cretaceous Symposium, Warsaw, Poland, pp. 249–250.

43. López-Horgue, M.A., Owen, H.G., Aranburu, A., Fernández-Mendiola, P.A., García-Mondéjar, J., 2009. Early late Albian (Cretaceous) of the central region of the Basque-Cantabrian basin, northern Spain: biostratigraphy based on ammonites and orbitolinids. Cretaceous Research 30, 385–400.

44. López-Horgue, M.A., Owen, H.G., Rodriguez-Lázaro, J., Orue-Etxebarria, X., Fernández- Mendiola, P.A., García-Mondéjar, J., 1999. The late Albian–lower Cenomanian stratigraphic succession near Estella-Lizarra (Navarra, central northern Spain) and its regional and interregional correlation. Cretaceous Research 20, 369–402.

45. Meek, F.B., 1876. Descriptions and illustrations of fossils from Vancouver’s and Sucia Islands, and other northwestern localities.

46. Morton, S.G., 1834. Synopsis of the Organic Remains of the Cretaceous Group of the United States: Illustrated by Nineteen Plates. To which is Added an Appendix, Containing a Tabular View of the Tertiary Fossils Hitherto Discovered in North America. Key \& Biddle.

47. Navarro-Ramirez, J., Bodin, S., Consorti, L., Immenhauser, A., 2017. Response of western South American epeiric-neritic ecosystem to middle Cretaceous Oceanic Anoxic Events. Cretaceous Research 75, 61–80.

48. Neumayr, V.H.M., 1875. Die Ammoniten der Kreide und die Systematik der Ammonitiden. Zeitschrift der Deutschen geologischen Gesellschaft XXVII, 854-942.

49. Neumayr, V.H.M., Uhlig, V., 1881. über Ammonitiden aus den Hilsbildungen Norddeutschlands. Palaeontographica (1846–1933), 129-203.

50. Owen, H.G., 2012. The Gault Group (Early Cretaceous, Albian) in East Kent, S. E. England; its lithology and ammonite biozonation. Proceedings of the GeologistśAssociation 123, 742–765.

51. Owen, H.G., Mutterlose, J., 2006. Late Albian ammonites from off-shore Suriname-implications for biostratigraphy and palaeobiogeography. Cretaceous Research 27, 717–727.

52. Parkinson, J., 1811. The Fossil Starfish, Echini, Shells, Insect, Amphibia, Mammalia &c. Organic remains of a former world an examination of the mineralized remains of the vegetables and nimals of the antediluvian world; generally termed extraneous fossils (3a ed.). C. Whittingham… and Published by J. Robson.

53. Parona, C.F., Bonarelli, G., 1896. Fossili Albiani d’ Escragnolles, del Nizzardo e della Liguria occidentale. Paleontographia italica: memorie di paleontologia II, 53-112.

54. Reboulet, S., Rawson, P.F., Moreno-Bedmar, J.A., Aguirre-Urreta, M.B., Barragán, R., Bogomolov, Y., Company, M., González-Arreola, C., Idakieva Stoyanova, V., Lukeneder, A., Matrion, B., Mitta, V., Randrianaly, H., Vasicek, Z., Baraboshkin, E.J., Bert, D., Bersac, S., Bogdanova, T.N., Bulot, L.G., Latil, J.-L., Mikhailova, I.A., Ropolo, P., Szives, O., 2011. Report on the 4th International Meeting of the IUGS Lower Cretaceous Ammonite Working Group, the “Kilian Group” (Dijon, France, 30th August 2010). Cretaceous Research 32, 786–793.

55. Renz, O., 1968. Über die Untergattungen *Venezoliceras* Spath und *Laraiceras* n . subgen . der Gattung *Oxytropidoceras* Stieler (Ammonoidea) aus den Venezolanischen Anden. Eclogae Geologicae Helvetiae 61, 615–655.

56. Renz, O., 1982. The Cretaceous Ammonites of Venezuela. Birkhäuser Verlag, Basel, Switzerland.

57. Royo y Gómez, J., 1942. Datos para la geología económica de Nariño y Alto Putumayo, Compilación de estudios geológicos oficiales en Colombia, pp. 53–260.

58. Saeid, E., Bakioglu, K.B., Kellogg, J., Leier, A., Martinez, J.A., Guerrero, E., 2017. Garzón Massif basement tectonics: Structural control on evolution of petroleum systems in upper Magdalena and Putumayo basins, Colombia. Marine and petroleum geology 88, 381–401.

59. Sarmiento-Rojas, L.F., 2019. Cretaceous Stratigraphy and Paleo-Facies Maps of Northwestern South America, in: Cediel, F., Shaw, R.P. (Eds.), Geology and Tectonics of Northwestern South America: The Pacific-Caribbean-Andean Junction. Springer International Publishing, Cham, pp. 673–747.

60. Spath, L.F., 1925. On Upper Albian Ammonoidea from Portuguese East Africa, with an appendix on Upper Cretaceous ammonites from Maputoland. Annals of the Transvaal Museum 11, 179–200.

61. Spath, L.F., 1934. A Monograph of the Ammonoidea of the Gault. Part XI. Monographs of the Palaeontographical Society 86, 443–496.

62. Spath, L.F., 1941. A Monograph of the Ammonoidea of the Gault. Part XIV. Pages 609–668; Plates LXV–LXXII. Monographs of the Palaeontographical Society 95, 609–668.

63. Stieler, V.C., 1920. Über sogenannte Mortoniceraten des Gault. Zentralblatt far Mineralogie, Geologie und Paláontologie, 345-352.

64. Tschopp, H.J., 1953. Oil Explorations in the Oriente of Ecuador, 1938-1950. AAPG bulletin 37, 2303-2347.

65. Villamil, T., 1998. Chronology relative sea level history and a new sequence stratigraphic model for basinal Cretaceous facies of Colombia. Society for Sedimentary Geology(SEMP), 161–216.

66. Villamil, T., Pindell, J.L., 1998. Mesozoic paleogeographic evolution of northern South America: Foundations for sequence stratigraphic studies in passive margin strata deposited during non- glacial times, in: Pindell, J.L., Drake, C. (Eds.). SEPM special publication, pp. 283–318.

67. Wiedmann, J., Owen, H.G., 2001. Late Albian ammonite biostratigraphy of the Kirchrode I borehole, Hannover, Germany. Palaeogeography, Palaeoclimatology, Palaeoecology 174, 161– 180.

68. Wright, C.W., 1952. A classification of the Cretaceous ammonites. Journal of Paleontology, 213-222.

69. Wright, C.W., Calloman, J.H., Howart, M.K., 1996. Cretaceous ammonoidea. Treatise on Invertebrate Paleontology, Part L, Mollusca 4, Revised. Geological Society of America, Boulder, Colorado, and University of Kansas Press, Lawrence, Kansas.

70. Young, K., 1958. *Graysonites*, a Cretaceous ammonite in Texas. Journal of Paleontology 32, 171–182.

